# The pore conformation of lymphocyte perforin

**DOI:** 10.1101/2021.07.03.450947

**Authors:** ME Ivanova, N Lukoyanova, S Malhotra, M Topf, JA Trapani, I Voskoboinik, HR Saibil

**Affiliations:** Institute of Structural and Molecular Biology, Birkbeck, University of London, Malet St, London WC1E 7HX, UK; Imperial College London, Hammersmith Campus, Du Cane Road, London W12 0NN, UK; Scientific computing department, Science and Technology Facilities Council, Rutherford Appleton Laboratory, Fermi Ave, Harwell, Didcot OX11 0QX, UK; Centre for Structural Systems Biology, Leibniz-Institut für Experimentelle Virologie and Universitätsklinikum Hamburg-Eppendorf (UKE), Hamburg, Germany; Peter MacCallum Cancer Centre, 305 Grattan St, Melbourne VIC 3000, Australia

## Abstract

Perforin is a pore-forming protein that facilitates rapid killing of pathogen-infected or cancerous cells by the immune system. Perforin is released from cytotoxic lymphocytes, together with pro-apoptotic granzymes, to bind to the plasma membrane of the target cell where it oligomerises and forms pores. The pores allow granzyme entry, which rapidly triggers the apoptotic death of the target cell. Here we present a 4 Å resolution cryo-EM structure of the perforin pore, revealing new inter- and intra-molecular interactions stabilising the pore. During assembly and pore formation, the helix-turn-helix motif at the bend in the central β-sheet moves away from the bend to form an intermolecular contact. Cryo-electron tomography shows that prepores form on the membrane surface with minimal conformational changes. Our findings suggest the sequence of conformational changes underlying oligomerisation and membrane insertion, and explain how several pathogenic mutations affect function.

## Introduction

Cytotoxic T lymphocytes (CTLs) and natural killer (NK) cells are essential for survival because they eliminate viral infection or destroy cancerous cells ^1,2^. In order to kill target cells, activated CTLs synthesise secretory vesicles (colloquially known as ‘lytic granules’) containing the pore-forming protein perforin ^3–5^ and pro-apoptotic serine proteases granzymes ^6^. CTLs form tight contacts, known as immune synapses, with virally-infected or transformed cells, and release the contents of lytic granules by exocytosis into the immune synapse ^7^. In the extracellular medium, which contains 1-1.3 mM of free Ca^2+^, perforin binds to the target cell plasma membrane where it oligomerises into arcs and rings, and transforms into pores that allow the entry of granzymes, triggering apoptosis ^8,9^.

Complete congenital loss of perforin function invariably results in Type 2 familial haemophagocytic lymphohistiocytosis (FHL) – a life-threatening autosomal recessive disorder that ultimately stems from cytokine hypersecretion and uncontrolled macrophage activation ^10^; if left untreated, the median survival of individuals who inherit two null perforin alleles is just 2 months ^11^. Hypomorphic perforin mutations are associated with atypical/late-onset FHL ^12^, lymphoma and other cancers ^13–16^.

Perforin is a member of the Membrane Attack Complex PerForin/Cholesterol Dependent Cytolysin (MACPF/CDC) superfamily of pore-forming proteins ^17–19^. Members of this superfamily are found in all kingdoms of life and are involved in diverse processes including toxic attack ^20^, immune defence ^21^, development ^22^, pathogen invasion ^23^ and inflammation ^24^. These proteins characteristically undergo a major conformational change to convert water-soluble monomers into oligomeric transmembrane pores. The conserved MACPF domain consists of a central, bent β-sheet and three helical regions: two α-helical bundles known as transmembrane hairpins (TMHs) 1 and 2, and a third helical bundle termed helix-turn-helix (HTH) motif. Although the core MACPF domain topology is conserved, the surrounding domains are highly divergent. Upon membrane docking, TMH 1 and 2 extend into amphipathic β-hairpins that assemble into a giant β-barrel spanning the target cell membrane ^25^. It is still unclear what signal triggers the transition between soluble and pore conformations, but it has been shown that prior to membrane insertion many MACPF/CDC proteins oligomerise into circular prepores – a pore precursor assembly that binds the membrane but is not inserted ^26^. It is not always necessary to assemble a complete ring, as arcs and incomplete rings have been shown to perforate the membranes *in vitro* ^27–29^, but the physiological relevance of these assemblies is not clear. Other members of the MACPF/CDC family such as the membrane attack complex (MAC) do not form a prepore and are able to add subunits after the initial complex has been inserted ^30,31^.

The structure of the soluble form of perforin was previously solved by X-ray crystallography ^32^. The perforin subunit consists of three domains: the conserved MACPF domain, the membrane docking C2 domain and the EGF-like domain (Figure 1A). Structures of several MACPF and MACPF-like proteins in the membrane-bound state, such as the MAC ^30^, the macrophage protein perforin-2/MPEG ^33^ and the more distantly related gasdermin A3 ^34^, have been determined by cryo-electron microscopy (cryo-EM) with near atomic resolution, but do not feature the perforin domains and do not host the sites of pathogenic mutations found in perforin. Here we present the structure of the perforin pore, revealing its unique domain movements and explaining why some of the clinically-relevant mutations are deleterious for function, including some centrally involved in the conformational transition.

**Figure 1.**
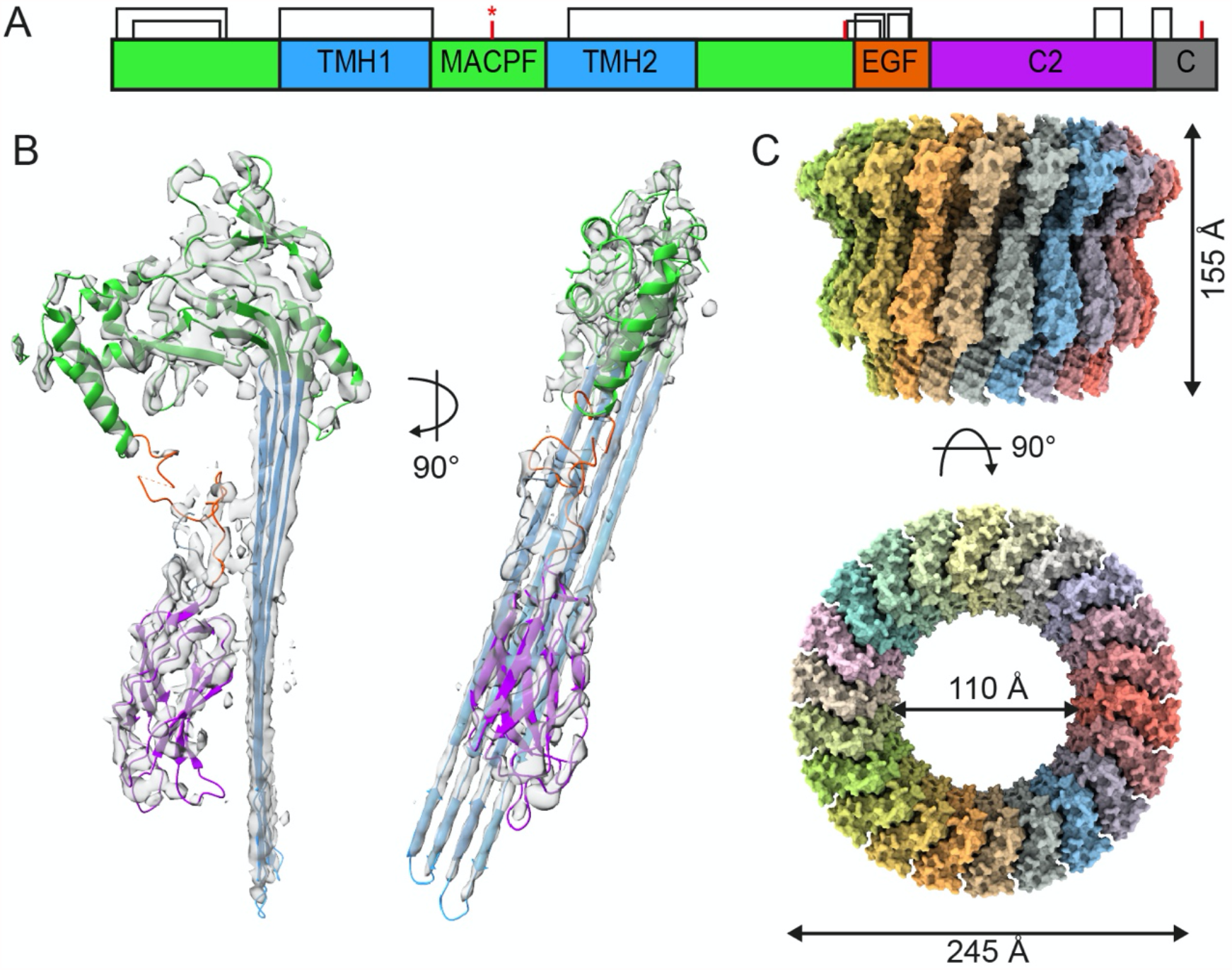
Overview of perforin structure. A. Domain structure of perforin. Disulphide bonds are indicated by black brackets, glycosylation sites by red bars, and N204 glycosylation site by a red asterisk. B. Fitting of the perforin model into the density corresponding to an inserted subunit. C. Overview of the perforin pore with C22 symmetry.

## Results

### Molecular architecture of a perforin pore

Murine wild-type perforin was recombinantly expressed and purified from baculovirus-infected insect cells as described previously ^35^. Perforin pores were assembled on phosphatidylcholine (PC) liposomes, in the presence of CaCl_2_ (see Materials and Methods). Detergent screening identified the commonly used detergent Triton X-100 as the best solubilising agent for perforin pores, as shown by negative stain EM (Supp. Fig. 1A). The detergent solubilised samples mainly contained complete perforin rings, suggesting that the incomplete rings, usually abundant in perforin pore preparations ^36^, are unstable in the presence of Triton-X-100.

We used single particle cryo-EM to solve the structure of the solubilised perforin pore. Inspection of two-dimensionally (2D) classified top views of perforin pores revealed particles with symmetry ranging from C15 to C26 with most having 21 to 23-fold symmetry (Supp. Fig. 2). Due to flexibility and size heterogeneity of the pores, particle subtraction followed by focused refinement was implemented to extract and resolve a wedge of the perforin pore, with a final resolution of 4.0 Å. The local resolution of the map ranged from 3.9 to 6.7 Å with the β-barrel being the best-defined feature (Supp. Fig. 1D and 1E). This map was used to build a molecular model of the perforin pore (Figure 1A).

The 22-fold perforin pore has a diameter of ∼245 Å and height of ∼155 Å (Figure 1B). The narrowest part of the pore has an opening of ∼110 Å in diameter which is sufficient to allow passage of a granzyme B monomer or a granzyme A dimer into the target cell. The MACPF, EGF-like and C2 domains of perforin are outside the pore, while the luminal side is composed of a giant β-barrel with diameter of ∼145 Å and height of ∼125 Å. An earlier study based on low-resolution data and labelling of sites in the perforin C-terminus suggested that the orientation of the perforin molecule in the pore is reversed relative to other members of the CDC/MACPF superfamily ^32^. Although that orientation has always seemed unlikely, no further experimental evidence on perforin has emerged to establish the orientation until now. The C-terminal tail of perforin is not ordered in our structure, and it has been previously shown that this region is very flexible and cleaved during maturation ^37,38^. The overall architecture of the perforin molecule is supported by nine disulphide bonds which are scattered over all domains (Figure 1A). Three perforin glycosylation sites have been previously reported: N204, N375 and N548 ^39^, but density for only N204 glycosylation is observed in the pore structure.

Each perforin subunit contributes four antiparallel β-strands towards assembly of the β-barrel. The bottom 30 Å of the β-barrel forms the transmembrane pore, with the rest of the perforin ring located above the lipid bilayer (Figure 2A). The β-strands enter the membrane at an angle of ∼18º to the pore axis, making the right-handed twist characteristic of β-barrels formed by L-amino acids. There is a two-residue shift in register between adjacent subunits (Supp. Figure 1E) giving a shear number of 44 for the 22-fold symmetric pore, which equals the total number of antiparallel β-strands in the circular structure. This agrees well with theoretical predictions of giant β-barrel architecture ^40,41^. The membrane-facing portion of the β-barrel is lined by mostly hydrophobic sidechains, whereas the luminal side is mostly hydrophilic (Supp. Fig 1F).

**Figure 2.**
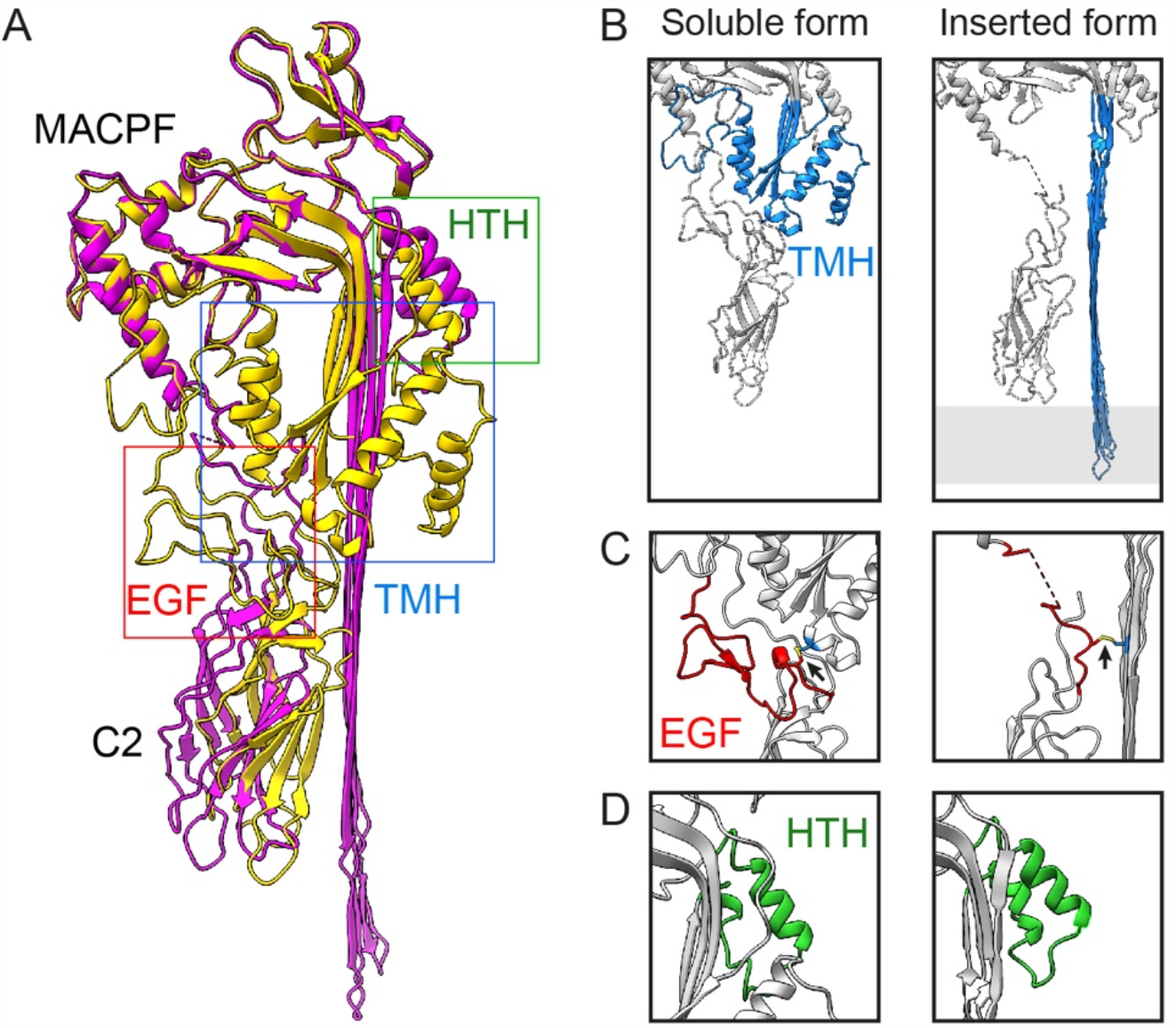
Conformational change of perforin upon insertion into the membrane. A. Superimposition of soluble (yellow, pdb code 3NSJ) and inserted (magenta) models of perforin. B, C and D. Structural comparison of TMH regions (B), EGF domain (C) and HTH motif (D) in soluble (left) and inserted (right) forms of perforin. In panel B the membrane position is indicated in grey. For panels B, C and D, domains are coloured according to the labels in A (HTH, green; TMH, blue; EGF, red) and the rest of the structure (inserted or soluble) is in grey. In C, the disulphide bond between the EGF and TMH is shown in yellow and indicated by arrows.

The narrowest part of the pore is formed by the helix-turn-helix (HTH) motif, which is enriched in negatively charged residues (Supp. Fig 1G). It has been previously shown that perforin pores show a preference for cationic cargos over neutral or negatively charged ones ^42^. In comparison, the Streptolysin O pore contains both positively and negatively charged amino acids in its HTH motif and is able to deliver a wide range of molecules irrespective of charge, although its wider diameter may lessen any charge effects ^43^ (Supp. Fig. 1G). A recent study of the gasdermin D pore shows that a charged lining in a pore of similar diameter to perforin confers substrate selectivity (Xia et al, 2021). In the perforin HTH motif, D308, D312 and E322 form a negatively charged ring on the luminal side of the pore may facilitate diffusion of granzyme B into the target cell. Notably, D312 is conserved in higher vertebrates and a mutation of this residue to valine (D313V in human perforin) has been found in a cancer patient ^44^. Human and murine perforin share 68% identity; the human protein has one additional residue at the cleaved signalling N-terminal peptide. While expression levels of this mutant perforin are comparable to those of the wild-type perforin, the lytic activity is reduced seven-fold in the context of mouse perforin.

### Conformational transition upon membrane insertion

Comparison of soluble and pore conformations reveals a major reorganisation of the protein upon membrane insertion (Figure 2). The upper part of the MACPF domain is quite rigid and remains largely unchanged; superposition of the soluble and inserted forms of the protein gave an r.m.s.d. of 1.65 Å over 189 Cα atoms in this region (Figure 2A, shown in green). The main conformational change happens in the lower part of the MACPF domain where helices comprising TMH 1 and 2 regions refold to form antiparallel membrane-spanning β-strands (Figure 2B). At the same time the central β-sheet unbends by 17º (Supplementary figure 3A). The EGF-like domain, which acts as a linker between the MACPF domain and the membrane-associated C2 domain, is covalently bound to TMH 2 through the disulphide bond between Cys407 and Cys241. Upon membrane insertion TMH 2 pulls the EGF domain 10 Å towards the newly assembled β-barrel (Figure 2C), so that the EGF domain fills in the space previously occupied by TMH 1. EGF-like domains are characterised by the presence of three or four canonical disulphide bonds ^45^, but the loops connecting conserved cysteines greatly vary in length and structure and can adopt a range of conformations, resulting in a very flexible architecture.

The helical arrangement of TMH 1 and 2 in the soluble form of the protein is stabilised by an extensive network of hydrogen bonds formed between this part of the molecule and adjacent domains: the upper part of MACPF domain, EGF-like domain and C-terminal tail (Supplementary Figure 3B). With the extrusion of TMH 1 and 2, the flexible EGF domain and C-terminal tail make few contacts and become extremely mobile, so that they could not be accurately fitted in the density. Therefore the structures of the EGF domain and C-terminus are less reliable than other parts of the molecule. Of the residues making interactions with TMH 1 and 2 in the soluble form of the protein, many are mutated in FHL patients (mutations R54C, H222R, H222Q, R361W, R410P, R410W in human numbering) ^2,44,46^. Two of these residues (H222 and R361) are conserved from fish to humans (Supp. Figure 4). Expression of the H222R mutant was shown to be undetectable or greatly reduced compared to the WT protein, suggesting this mutation leads to protein misfolding, whereas the mouse forms of H222Q and R361W variants were expressed in rat basophilic leukaemia (RBL) cells at levels equivalent to WT perforin ^47,48^. RBL cells transfected with H222Q mutant mouse protein had no detectable cytotoxic activity ^47^ suggesting this mutation affects the function of the protein rather than its stability. The structure of mouse perforin in its soluble form shows that the side chain of the equivalent residue (H221) makes a hydrogen bond with the main chain oxygen of W128, which is located in the loop between the two helices in TMH1. Substitution of histidine by (similarly sized) glutamine does not perturb the overall architecture of the molecule, but allows the formation of a stronger hydrogen bond with W128, which in turn disfavours the conformational transition required for membrane insertion of the mutant molecule. The loss of pore forming activity confirms that H222 is important for perforin function in disrupting the target cell membrane.

Another significant conformational change happens on the luminal side of the pore. The second helix in the third helical bundle, which is unfolded in the soluble monomer, assembles to form three helical turns completing the helix-turn-helix (HTH) motif facing the lumen of the pore at the top of the β-barrel (Figure 2D). The HTH moves up slightly and tilts by 6º away from the bend of the central β-sheet, a displacement that was previously proposed to unlock the conformational change needed for membrane insertion ^49^. This movement is supported by the movement of the side chain of R298 which makes a new interaction with the bottom loop of the HTH motif (Supplementary Figure 3C). In the soluble form of perforin, R298 makes a hydrogen bond interaction with the mainchain oxygens of H292 and Y295 stabilising the turn between the central β-sheet and helices of TMH2. R298 is absolutely conserved from fish to humans (Supp. Figure 4). This residue has been found to be mutated in FHL patients and this mutation has a detrimental effect on perforin expression and NK cell function ^44,47^.

### Oligomerisation interface

The primary oligomerisation interface is located on the flat face of the globular MACPF domain of perforin. It has been previously shown that E343 of one subunit interacts with R213 on the adjacent monomer ^32^, and that a salt bridge linking the two residues is indispensable for lytic function ^50^. The structure of the inserted form of perforin shows that these two residues are located in close proximity to each other with Arg and Glu side chains pointing towards each other (Figure 3A). Reversing the charge of either one of these residues causes complete loss of function, but reversing both of them restores WT function ^50^. A number of other charged residues are located on the flat surface of the MACPF domain and are likely to contribute towards the overall intermolecular affinity.

**Figure 3.**
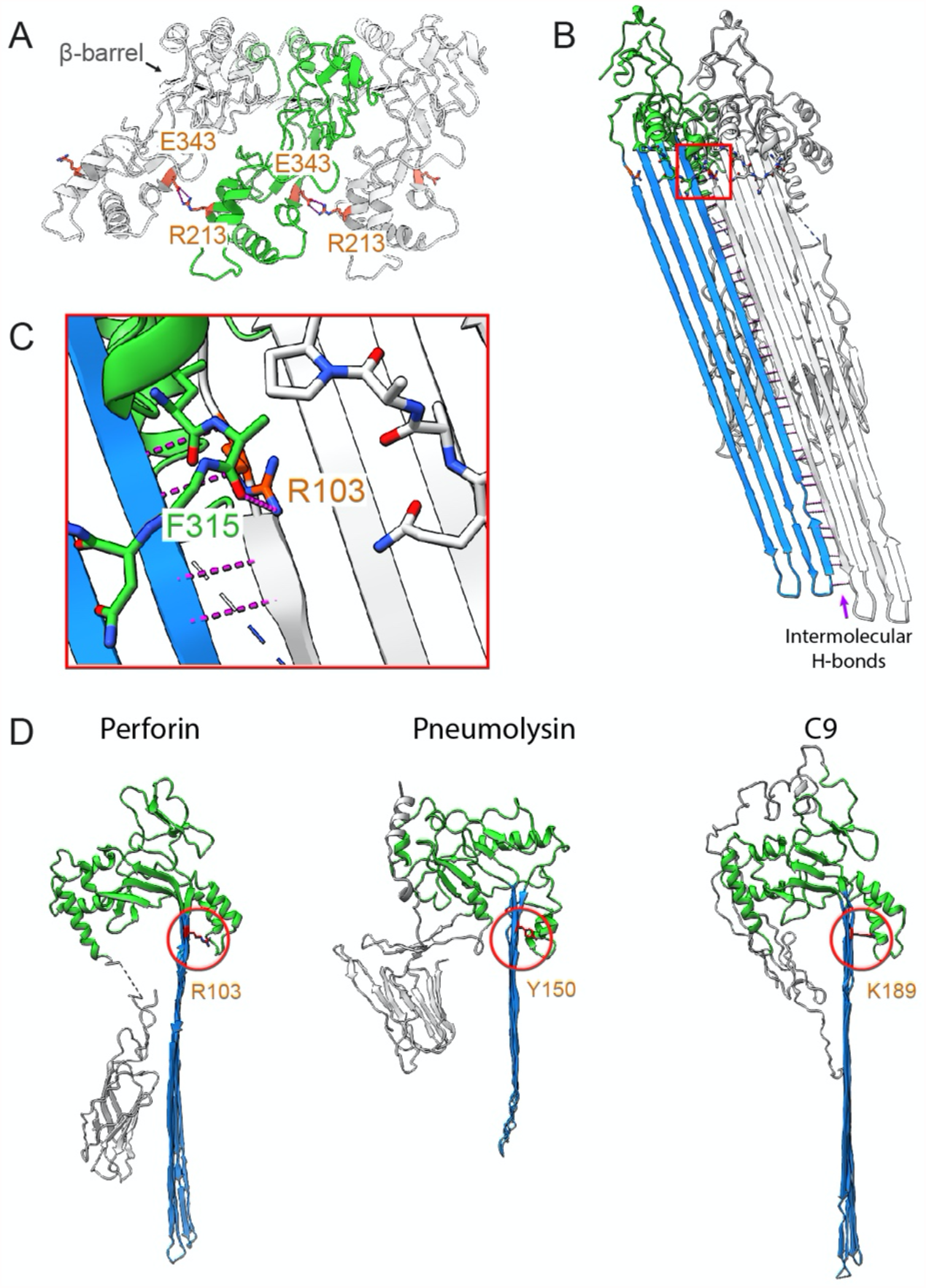
Oligomerisation interface. A. Top view of the perforin oligomer; the central perforin subunit is highlighted in green, with residues making intermolecular interactions in orange. B. Oligomerisation interface in the β-barrel with one subunit shown in colour and the adjacent one in grey. Intermolecular hydrogen bonds are shown in pink. C. Secondary oligomerisation interface formed between HTH motif of one subunit (shown in colour) and Arg103 in the β-barrel of the next (shown in light grey, Arg103 is highlighted in orange). This region is highlighted with a red box in panel C. D. Comparison of perforin with other MACPF proteins with the residues equivalent to Arg103 of perforin (Tyr150 and Lys189 of pneumolysin and C9 respectively) circled in red and shown in red sticks.

The second oligomerisation interface is formed by the HTH motif, mentioned above, on the luminal side of the pore. The HTH motif is formed of an insertion at the bend of the central β-sheet and it must be displaced away from the core of the MACPF domain for the TMH2 helices to refold into β-hairpins. Once displaced, it adopts a new conformation that is supported by the hydrogen bond formed between the side chain of R103 from a β-hairpin of one subunit and the mainchain oxygen of F315 located at the HTH turn of the adjacent subunit (Figure 3B). Comparison with other members of the MACPF family showed that this interaction is conserved between different pore forming proteins with a tyrosine or a lysine located at equivalent positions at the top of the first β-hairpin (Figure 3C).

Finally, once TMH 1 and 2 transition from α-helices to β-strands, the third, and strongest, oligomerisation interface is formed by β-hairpins from adjacent monomers. Antiparallel β-strands form the β-barrel via an extensive network of hydrogen bonds between the main-chain atoms of neighbouring strands (Figure 3D). Giant β-barrels can adopt variable curvature, allowing the pores to be built with a wide range of symmetries and pore diameters.

### Cryo-electron tomography of perforin prepores

We previously showed that prior to membrane insertion perforin molecules can form prepores – oligomers docked on but not inserted into membranes, typically consisting of up to eight subunits ^36^. They are initially loosely packed, but become more ordered over time. To characterise the perforin conformation in these assemblies, we used a TMH1-lock mutant of perforin (A144C-W373C), in which TMH1 is tethered to the core of the MACPF domain ^36^. This prevents the refolding of either TMH region into a β-hairpin and thereby completely blocks membrane insertion. The engineered disulphide bond in this mutant tethers the TMH1 region to a conserved MACPF α-helix. Perforin prepores are less ordered than mature pores and could not be isolated from liposomes, so it was necessary to use cryo-tomography to study the prepores on liposomes. For prepore formation, liposomes containing 10% biotinylated lipids were immobilised on functionalised EM grids containing streptavidin, in order to prevent liposome aggregation upon addition of perforin in the presence of Ca^2+^.

Multiple perforin assemblies, mainly incomplete rings, were clearly visible on the surface of the liposomes (Figure 4A). The height of these assemblies, 10.5 - 11 nm, agrees well with prepore height measurements by atomic force microscopy ^36^. The shape of the perforin molecule crystal structure is easily recognisable in the density sections, with the narrower Ca^2+^-binding C2 domain docked on the membrane surface (Figure 4B). The crystal structure of the soluble perforin monomer can be manually docked into the prepore density, qualitatively demonstrating that no major conformational change is required for prepore formation (Figure 4C).

**Figure 4.**
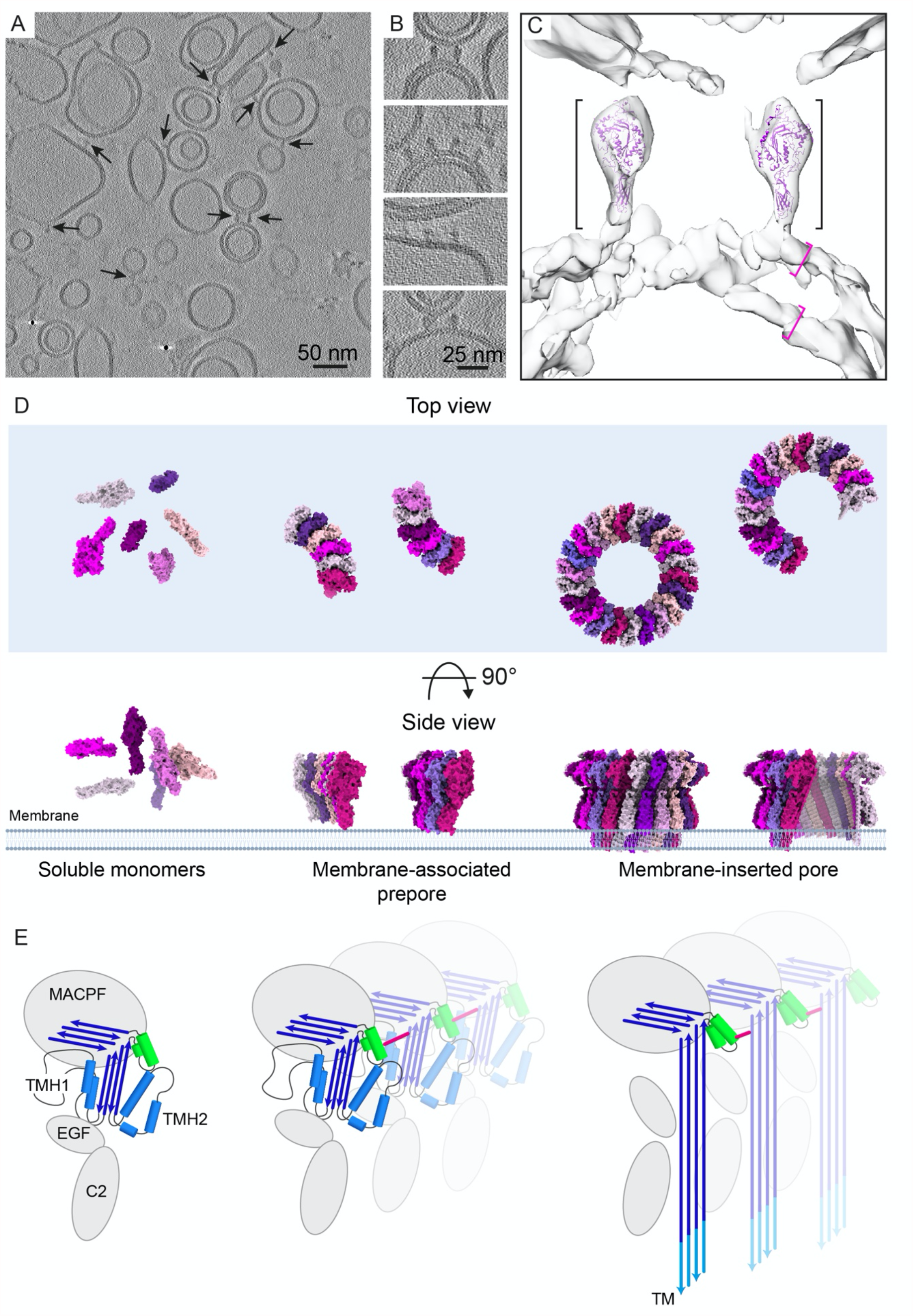
Cryo-tomography of perforin prepores. A. Overview of a typical cryo tomogram of liposomes with attached perforin prepores. Prepore assemblies are highlighted with arrows. B. Close up view of prepores attached to the lipid bilayers. C. Manual docking of soluble perforin monomer (purple, PDB code 3NSJ) into the 3D volume of a perforin prepore on a double shelled liposome. Density corresponding to the prepore is indicated by black brackets and density corresponding to each membrane bilayer is marked by pink brackets. D. Model of the molecular assemblies during pore formation. E. Schematic representation of the conformational changes and assembly interactions. The HTH region is shown in green, the bond between the HTH and a neighbouring β-strand in pink, the initial β-sheets in blue and the transmembrane region in cyan.

## Discussion

In this study we have used single-particle cryo-EM to determine the structure of membrane-inserted perforin pore at near-atomic resolution. In combination with a previously solved crystal structure of the soluble perforin monomer, this work describes the conformational changes that lead to membrane insertion of perforin.

Perforin is the key effector of cytotoxicity inflicted by activated CD8+ T cells and NK cells, mediated by its ability to form transmembrane pores that deliver granzymes to the target cell cytosol. Various human disorders have been associated with dysregulated biosynthesis or mutations that affect the perforin structure. These mutations are spread throughout the MACPF and C2 domains of the protein ^12,51^ (Supp. Fig. 4 and 5, all numbering used in this section corresponds to the human perforin sequence for comparison with clinical data). The most serious of these disorders is type 2 FHL (FHL2), which is specifically associated with loss of perforin expression or function ^12^. Current treatment of FHL includes chemotherapy, immunosuppression with high dose corticosteroids and antibiotic and/or antiviral drugs to address any potential infectious trigger. There is no treatment available to date that aims to restore perforin function or its delivery into the immunological synapse, although even mild changes in perforin activity might contribute to immune dysregulation ^52^. The structure of the inserted form of perforin provides insight in the mechanism of pore formation and suggests how some FHL-causing mutations affect protein structure and function.

The MACPF family includes hundreds of diverse proteins, but few have been extensively studied and structural information is available for even fewer. Medium resolution structures of the pneumolysin pore and the MAC complex in its inserted form have been determined by cryo-EM ^29,30,53^. The MACPF domains of these proteins are structurally conserved and contain a signature motif (Y/W)G(T/S)H(F/Y)X_6_GG, but otherwise share very little similarity and contain diverse auxiliary domains. The perforin domains play distinct roles: the MACPF domain is responsible for oligomerisation and membrane insertion, the C2 domain binds Ca^2+^ ions and docks the molecule onto the membrane, while the EGF domain provides flexibility, links the MACPF and C2 domains and is tethered to TMH2. Canonical EGF domains have diverse functions, including extracellular and intracellular signalling, ligand recognition, and mediating protein interactions. MAC proteins also contain EGF domains, but their position relative to the MACPF domain is different to the one in perforin. Moreover, EGF domains in MAC proteins lack a disulphide bond linking them to the TMH2 domain. This suggests that the EGF domains play different roles in different MACPF proteins.

Soluble perforin is monomeric ^50^ whereas in the presence of a lipid bilayer, perforin molecules dock onto the membrane surface via the Ca^2+^ binding sites at the base of the C2 domain and then oligomerise. Leung et al identified two possible types of perforin prepores, early and late prepores, with more compact subunit packing at the later stage ^36^. Both types of prepore formed shorter segments, with 2-5x fewer subunits than the pores. Pore formation could be induced by addition of reducing agent, to release the TMH regions. Prepore oligomerisation on the membrane is not reversible, ^36^, suggesting that oligomerisation involves a conformational change preceding β-barrel formation and membrane insertion. We speculate that the newly identified interaction between the HTH motif of one molecule and a β-strand of the adjacent subunit forms at the prepore stage and might stabilise the prepore and promote further assembly (Figure 4D). Displacement of the HTH domain away from the core of the MACPF domain is accompanied by formation of a new intermolecular interaction between the HTH motif and R298, which in turn slightly unbends the central β-sheet, releasing the TMH 2 helices for unfolding and conversion into β-hairpins. The HTH motif is located adjacent to the bend in the central β-sheet, flanked by two glycine residues that are conserved throughout the MACPF/CDC superfamily, allowing for flexibility at the bend in the β-sheet. Partial straightening of this bend is crucial to the MACPF/CDC conformational change. Our discovery of an intersubunit contact involving this region is highly significant, since it links triggering of the conformational change to oligomerisation (Figure 4E).

This study provides important insights into the structure of the clinically significant protein perforin in its membrane inserted form, as well as explaining the mode of action of some pathogenic mutations. Furthermore, our observations help us to understand the mechanism of pore formation by MACPF proteins and suggest the order of conformational changes required for membrane insertion.

## Methods

### Protein expression and sample preparation for electron microscopy analysis

Wild-type and TMH1-lock mutant (A144C-W373C) perforin were expressed in a baculovirus/insect cell system and purified from the supernatant as described by Voskoboinik and colleagues ^35,36^.

100 μL of phosphatidylcholine lipid dissolved in chloroform at 10 mg/mL (Avanti Polar Lipid, USA) was dried under nitrogen gas and resuspended in 1 mL of buffer containing 20 mM HEPES at pH 7.5, 150 mM NaCl and 5 mM CaCl_2_. Rehydrated lipid solution was sonicated in a water bath at 40°C for 10 min followed by flash freezing in liquid nitrogen. Thawing and sonication followed by flash freezing were repeated 3 times, yielding a solution of large multilamellar vesicles which was forced through a polycarbonate filter with 80 nm pore membrane around twenty times using the Avanti mini-extruder at 40°C. The resulting liposome suspension was stored at 4°C and used within 48 hours.

Perforin pores were assembled on liposomes by incubating 5 μL of the liposome solution with 15 μL of purified protein at concentration of 200-300 ng/ml for 15 min at 37°C. For the preparation of perforin pore complexes, the proteoliposomes were solubilized at a final concentration of 1% Triton X-100 at room temperature overnight. Solubilised pores were stored at 4°C and used within 48 hours.

### Negative-stain and cryo-electron microscopy of perforin pores

For negative-stain EM, 3 μL of solubilised perforin pores was applied onto a continuous carbon TEM grid (Electron Microscope Sciences, USA) freshly negatively glow discharged using PELCO Easiglow (Ted Pella, USA) and stained with 2% (wt/vol) uranyl acetate. Negatively stained specimens were examined with a T12 electron microscope (FEI, The Netherlands) operated at an acceleration voltage of 120 keV. Images were recorded with a 4kx4k UltraScan CCD camera (Gatan, USA) at a nominal magnification of 25,000–45,000 and ∼1.0–2.0 μm underfocus.

For cryo-EM UltrAuFoil R 2.0/2.0 grids (Quantifoil, Germany) were negatively glow discharged using a PELCO Easiglow and coated with graphene oxide as described in Cheng et al., 2020 ^54^. In order to increase particle concentration, 3 μL of the sample were applied twice with a 30 sec interval and 0.5 sec blotting prior to the second application inside the chamber of VItrobot Mark IV (Thermo Fisher Scientific, USA) at 4°C and 92 % humidity. Samples were vitrified in liquid ethane. Cryo EM data were collected at the ISMB Birkbeck EM facility using a Titan Krios microscope (Thermo Fisher Scientific, USA) operated at 300 keV and equipped with a BioQuantum energy filter (Gatan, USA) using a slit width of 20 eV. The images were collected with a post-GIF K3 direct electron detector (Gatan, USA) operated in super resolution mode, at a magnification of 64,000, corresponding to a physical pixel size of 1.34 Å. The dose rate was set to 16 e per pixel per second, and a total dose of 49.6 e per Å^2^ was fractionated over 50 frames. Data were collected using EPU software (Thermo Fisher Scientific, USA) with a defocus range of 1.5 μm - 3.3 μm. A total of 19,627 movies were collected. To mitigate preferred orientation of perforin pores on the continuous substrate, the microscope stage was tilted by -30º during data collection.

### Image processing

Electron micrograph movie frames were aligned by MotionCor2 ^55^ using the RELION v3.1 implementation ^56^. Super-resolution movies were additionally down sampled by a factor of 2, applied by Fourier binning within MotionCor2. All aligned movie frames were subsequently averaged into dose-weighted and non-weighted sums for further processing. Contrast transfer function (CTF) estimation of whole non-dose-weighted micrographs was initially performed with CTFFIND4 ^57^.

Particle coordinates were determined using Gautomatch (developed by Kai Zhang; https://www.mrc-lmb.cam.ac.uk/kzhang/Gautomatch/) and particles were extracted with a box size of 320 pixels using RELION. Extracted particles were imported into cryoSPARC v2 ^58^ for 5 rounds of 2D classification. Particles corresponding to 2D class averages showing clear features and having similar diameter were selected and all further processing was performed in RELION v3.1 unless otherwise stated (Suppl. Fig 2).

Particles were subjected to two rounds of 3D classification followed by one round of consensus 3D refinement. Particles corresponding to the class showing the most defined features and the most complete ring were selected for further processing. This was followed by two rounds of per-particle CTF estimation and further 3D refinement. These alignments served as the starting point for tracking beam-induced movement of individual particles, which was corrected using particle polishing in RELION. The final subset of particles included images of perforin pores containing mainly 22 monomers, but due to pore flexibility, the structure was refined without imposing symmetry; this yielded a reconstruction with 7.1 Å global resolution, estimated using the gold standard Fourier Shell Correlation (FSC) with a 0.143 threshold. Subsequently symmetry expansion was implemented such that each particle was assigned 22 orientations that corresponded to its symmetry-related views. These particles were then subjected to one round of 3D refinement with only local angular searches and local resolution estimation of the final map had been performed. A mask was prepared to only include five monomers of perforin that were best defined, while the rest of the pore was subtracted from the original images. Focused 3D refinement was performed to improve the resolution of this 5-subunit wedge of the perforin ring; this was followed by another round of 3D refinement with only local angular searches, yielding 4.1 Å resolution. To limit anisotropy of the map, ∼300,000 particles out of 5,000,000 symmetry -expanded copies were selected for the final refinement using rlnMaxValueProbDistribution criteria ^59^; the remaining particles, corresponding to over-represented views, were excluded. This procedure reduced the average resolution of the map from 4.1 Å to 4.5 Å, but improved the definition of structural features. A calibrated pixel size of 1.34 Å was applied for RELION postprocessing, yielding a global resolution of 4.0 Å, determined by gold standard Fourier shell correlation (FSC) with 0.143 threshold (Supp. Fig 1C). Local resolution was estimated in RELION using a windowed FSC_0.143_.

### Model building and analysis

For model building the final map was modified using Phenix Resolve software ^60^. The initial model was generated in Coot ^61^ using the previously determined crystal structure of a soluble perforin monomer ^32^ and an atomic model of membrane attack complex C9 protein built into a cryo-EM map ^30^. Model building was performed using a combination of Flex-EM (with disulfide restraints) ^62^, Coot and real-space refinement in Phenix ^63^. Model validation was performed using MolProbity ^64^ and the CCP-EM software suite ^65^. The pore model with 22 subunits was generated in UCSF Chimera ^66^. Density maps and models were visualised and figures were drawn in UCSF ChimeraX ^67^. Coulombic potentials of interaction interfaces were calculated and visualized in UCSF Chimera.

### Cryo-electron tomography of perforin prepores

Liposomes were prepared as described above using phosphatidylcholine (PC), cholesterol and 1,2-dipalmitoyl-sn-glycero-3-phosphoethanolamine-N-(biotinyl) (Biotinyl PE) at 6:3:1 molar ratio. Liposome concentration was checked by negative stain EM prior to sample preparation. To prevent liposome aggregation upon prepore formation we used C-SMART Streptavidin BioGrids (Dunes Sciences Inc., USA) with a thin layer of continuous carbon derivatised with streptavidin (http://www.dunesciences.com/biogrids.php) over a lacey carbon substrate. Streptavidin BioGrids were first rinsed with buffer containing 20 mM HEPES at pH 7.5, 150 mM NaCl and 5 mM CaCl_2_ by floating on a droplet, then incubated with diluted liposomes inside a humid chamber at 37°C for 10-20 min and rinsed on a buffer droplet again without blotting. Then 3 uL of TMH1-lock mutant (A144C-W373C) perforin diluted in the same buffer were added directly onto a grid at approximate molar ratio of 1:10,000 protein to lipid. Grid was then transferred into the chamber of a Vitrobot Mark IV and after 5-10 min incubation at 37°C and 90 % humidity, blotted and vitrified in liquid ethane. Immediately before blotting, 1 μl of 6 nm Protein A gold fiducials (Electron Microscope Sciences, USA) diluted 5 times with the above buffer were added directly onto the grid.

Tilt series were collected at the eBIC National facility using a Titan Krios microscope operated at 300 keV with a post-GIF K2 Summit direct electron detector (Gatan, USA) operating in counting mode, at a nominal magnification of 81,000, corresponding to a pixel size of 1.77 Å. The dose rate was set to 6.2 e per pixel per second. Tomo 3 software (Thermo Fisher Scientific, USA) was used to collect tilt series between −45° and 45° using a linear tilt scheme with 3° increments starting at 0° tilt with tracking before and after. A total exposure of ∼50 e/Å^2^ was fractionated over 93 frames with dose 1.6 e/Å^2^ dose per tilt. An energy slit with 20 eV width was used during data collection. A volta phase plate (VPP) was used to enhance the contrast and data collected with a nominal defocus range from 60 nm - 120 nm. The VPP was advanced to a new position for every tilt series with 40 sec activation time and total dose on the VPP ∼50 nC.

MotionCor2 v1.0.5 ^55^ was used for subframe alignment and dose weighting. Tilt series were aligned and reconstructed by weighted back-projection using IMOD/etomo ^68^.

## Data availability

The cryo-EM map and corresponding atomic model have been deposited in public repositories with accession codes EMD-XXX and PDB XXX respectively.

## Author contributions

M.E.I., N.L., J.A.T., I.V. and H.R.S. designed the experiments. M.E.I., N.L. and I.V. collected and analysed the data. M.E.I., S.M. and M.T. built and refined the atomic model. M.E.I. and H.R.S. wrote the manuscript with input from other authors.

## Acknowledgements

This work was funded by ERC grant 294408 and Wellcome Trust grant 106249/Z/14/Z to H.R.S, and Wellcome Trust (209250/Z/17/Z and 208398/Z/17/Z) to M.T.. Most of the cryo-EM data for this investigation were collected at the ISMB EM facility at Birkbeck College, University of London with financial support from Wellcome Trust (202679/Z/16/Z and 206166/Z/17/Z). We thank Diamond Light Source for access to the cryo-EM facilities at the UK National electron bio-imaging centre (eBIC, proposal EM14704) funded by the Wellcome Trust, the Medical Research Council UK and the Biotechnology and Biological Sciences Research Council. We thank Annette Ciccone and Sandra Verschoor for perforin preparation, David Houldershaw for IT support and script writing, Dan Clare for help with tomography data collection and Bart Hoogenboom for comments on the manuscript.

## Notes

### Competing Interest Statement

The authors have declared no competing interest.

