## Supplementary table and figures for "The pore conformation of lymphocyte perforin"

**Supplementary Table 1 Cryo-EM data collection, refinement and validation statistics**

| | |
|---|---|
| **Data collection and processing** | |
| Voltage | 300 |
| Electron exposure ($e^{-}$/Å$^2$) | 49.6 |
| Defocus range (μm) | 1.5 – 3.3 |
| Nominal magnification | 105000x |
| Super-resolution pixel size (Å) | 0.69 |
| Initial particle images (no.) | 1,062,243 |
| Final particle images (no.) | 229,789 |
| Map resolution (Å) (FSC = 0.143) | 4.0 |
| Map resolution range (Å) | 3.9 – 7.3 |
| **Model refinement** | |
| Initial model used (PDB code) | 3nsj |
| Model resolution (Å) (FSC = 0.5) | 7.27 |
| Model resolution (Å) (FSC = 0.143) | 3.96 |
| Map sharpening B factor (Å$^2$) | -95 |
| Model composition | |
| Non-hydrogen atoms | 4003 |
| Protein residues | 509 |
| Ligands | |
| Ca | 3 |
| NAG | 1 |
| B factors (Å$^2$) | |
| Protein | 129.7 |
| Ligand | 181.6 |
| R. m. s. deviations | |
| Bond lengths (Å) | 0.27 |
| Bond angles (°) | 0.55 |
| **Validation** | |
| MolProbity score | 1.99 (77$^{th}$ percentile*) |
| Clashscore | 11 (67$^{th}$ percentile*) |
| Poor rotamers (%) | 0 |
| Ramachandran plot | |
| Favoured (%) | 94 |
| Allowed (%) | 6 |
| Disallowed (%) | 0 |

*100th is the best among the structures, 0th is the worst.

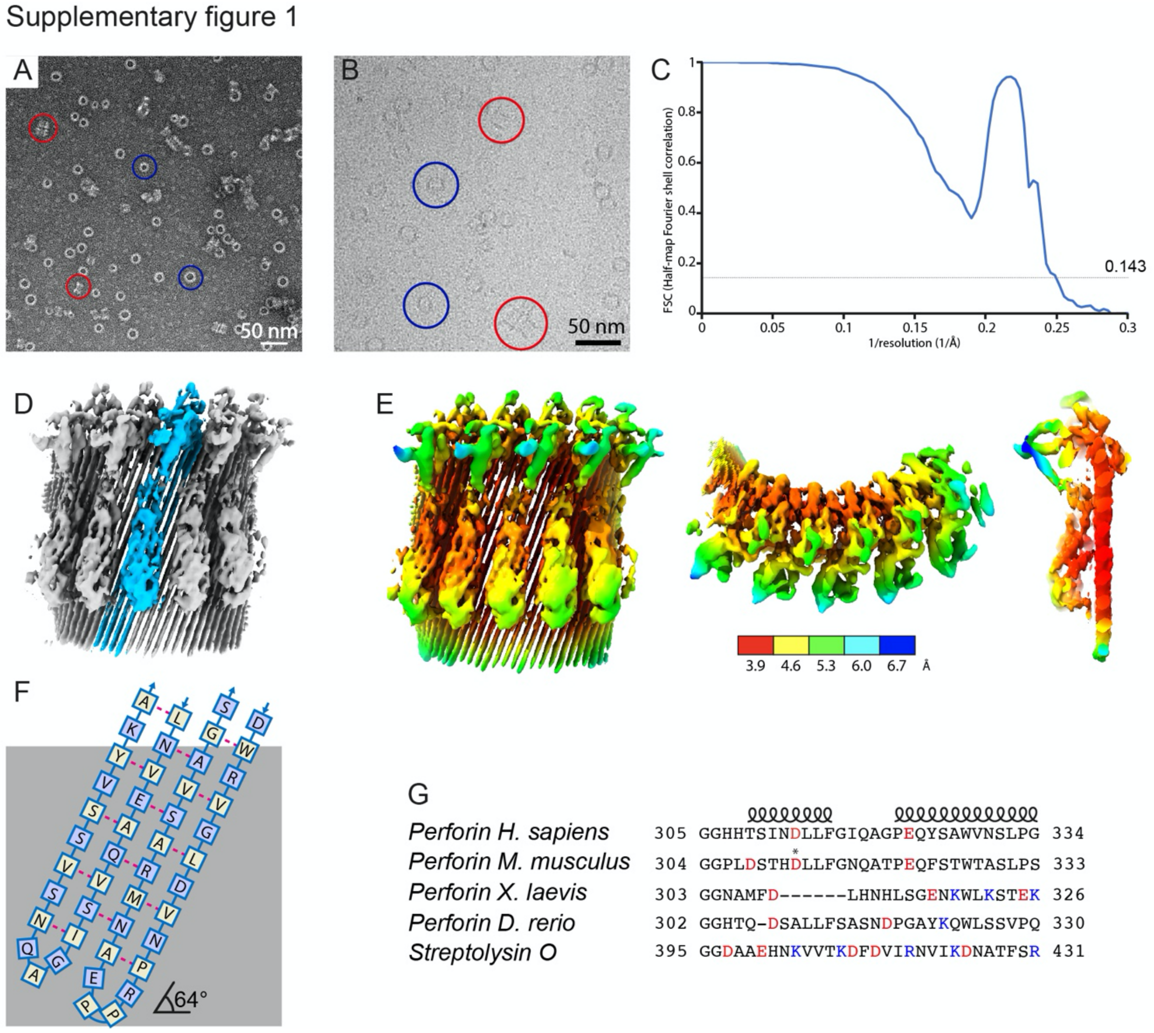

**Supplementary figure 1.**

A. Representative negative stain micrograph of perforin pores; end views are circled in blue, side views in red. B. Representative micrograph of vitrified perforin pores with example views circles as in A. C. Gold-standard Fourier shell correlation (FSC) curve calculated from two independently refined half-maps indicate an overall resolution of 4.0 Å at FSC = 0.143. There is a strong peak at $(4.8\ \text{Å})^{-1}$ because the highly ordered β-barrel is the dominant structural element. D. Overview of the obtained map of the wedge of a perforin pore. A single central perforin subunit is highlighted in blue. E. Local resolution estimate shown on front (left), top (middle) and cross-section (right) views of the cryo-EM density map reveals an average resolution of 3.9 Å or better in the β-barrel of the perforin pore. F. Arrangement of intramolecular hydrogen bonds (dashed red lines) formed within the transmembrane part (grey) of the β-barrel. Residues with sidechains pointing towards the membrane are shown in white squares, residues with sidechains pointing towards the lumen of the pore are shown in blue squares. G. Sequence alignment of HTH region of vertebrate perforins and the CDC protein Streptolysin O. Negatively charged residues are highlighted in red, positively charged residues in blue. D312 mentioned in the text is highlighted with an asterisk.

Supplementary figure 2

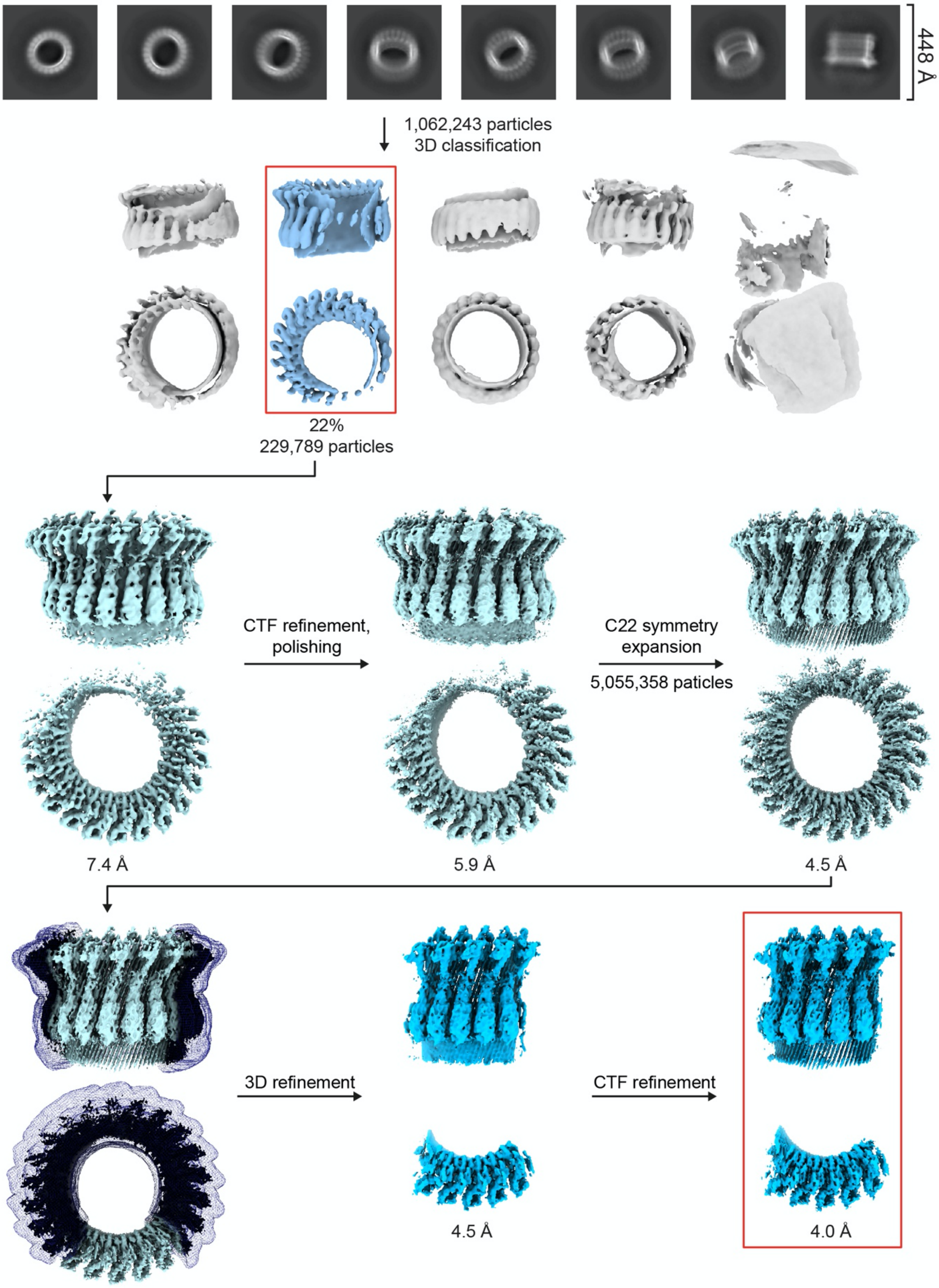

**Supplementary Figure 2. Structure determination pipeline of perforin pore**

Supplementary figure 3

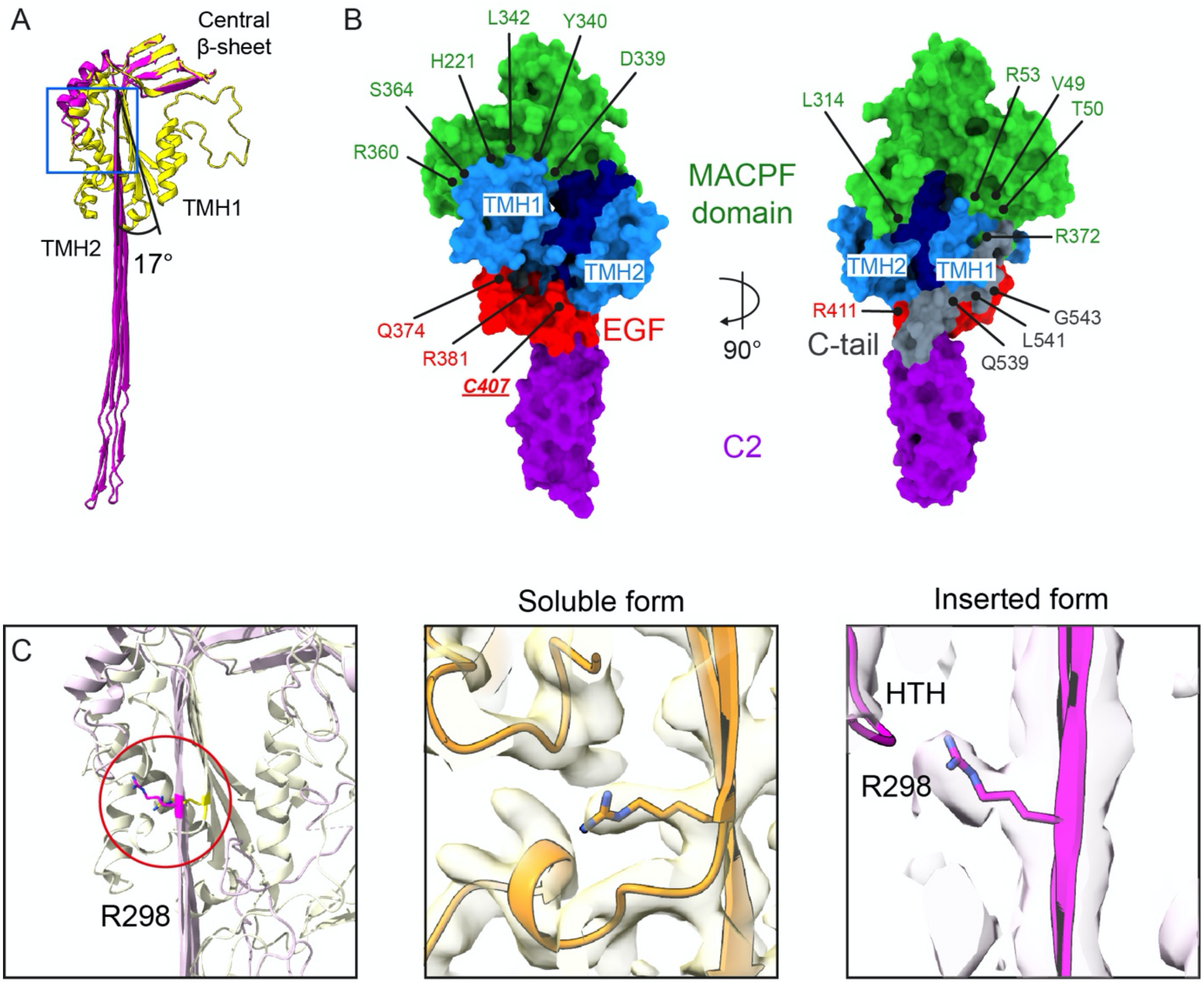

**Supplementary Figure 3**. **Intramolecular interactions in perforin**.
A. Superposition of the central β-sheet and TMH regions of soluble (yellow) and inserted (magenta) models of perforin showing the 17° unbending of the sheet. B. Residues supporting the helical arrangement of TMH 1 and 2 in the soluble form of perforin. Cysteine 407 (underlined) makes a disulphide bond with a cysteine in TMH 2. C. The conformational change of the R298 side chain supports the HTH motif movement. Left panel shows superposition of soluble (yellow) and inserted (magenta) forms of perforin, middle and right panel show the closeups of R298 (stick models) in the two maps.

Supplementary figure 4

```
                           11          21        31        41        51        61        71        81
Homo sapiens    MAARL-LLLGILL--LLLPLPVPAPCHTAARSECKRSHKFVPGAWLAGEGVDVTSLRRSGSFPVDTQRFLRPDGTCTLCENALQ
Mus musculus    MATCL-FLLGLFL---LLPRPVPAPCYTATRSECKQKHKFVPGVWMAGEGMDVTTLRRSGSFPVNTQRFLRPDRTCTLCKNSLM
Xenopus laevis  MLHFFMIICFIPLICSSAPLLHTVTCKQGTKTQCLK-ASFVPGHNLPARGIDIVTMKSAGNILFNTQQYMK-NNTCKLCQDSQD
Danio rerio     MVPVA-LLCSVLL--LYLTPPTSPACFLGKQAECQD-ASFVPGSDLAGEGFDITKMQRLGTFVFDMSKWDLNNNTCNLCKNPFM
                V                             C         MH       E CM   C               Y  R

         91        101       111       121       131       141       151       161       171
EGTLQRLPLALTNWRAQGSGCQRHVTRAKVSSTEAVARDAARSIRNDWKVGLDVTPKPTSNVHVSVAGSHSQAANFAAQKTHQDQYSFSTDTVECRFYS
RDATQRLPVAITHWRPHSSHCQRNVAAAKVHSTEGVAREAAANINNDWRVGLDVNPRPEANMRASVAGSHSKVANFAAEKTYQDQYNFNSDTVECRMYS
KDKWWKLPRALEQWNMQS-SCRRTMVSKVLQSPVLVAKEVASRVQNDWTNDLNLTCT-ELQGNVAMAGAQSKMVQFAFEKLHLDKYTFLSHKFECEYYS
QNEKQKLPVSAVYWRASQ-KCTMSLSSSVYESSESLVSSSTSSIENNWKTGLGIGNM-LSKVSLMVAGTNSRLSDYSMSKTKKDKFSFIKQSASCVYYS
       T V   R      F                W   H  C                      S       V          N R      C

181       191       201       211       221       231       241       251       261       271
FHVVHTPPLHPDFKRALGDLPHHFNASTQPAYLRLISNYGTHFIRAVELGGRISALTALRTCELALEGLTDNEVEDCLTVEAQVNIGIHGSISAEAKAC
FRLVQKPPLHLDFKKALRALPRNFNSSTEHAYHRLISSYGTHFITAVDLAGRISVLTALRTCQLTLNGLTADEVGDCLNVEAQVSIGAQASVSSEYKAC
FQMSRHWNLSPHFLKALVQLPNYYNSRSNVQYQQLIASYGTHYITDATMGGQLQEVTALRTCEVALSDSTMQLVKDCREAEIMAAVVGPNNT-MLSQIC
YRIASSAPLHHELRDDFSNLPATYDNQTKESYLSLVEKFGTHYIIQVKLGGGVQSVTSVKQCMATLQDLSVDEVKTCLDVEASASVMGRISVDTAYRHC
  G         L    R  T             I    SIR DW      H       G           S        KD                Y

 281       291       301       311       321       331       341       351       361       371
EEKKKKHKMTASFHQTYRERHSEVVGGHHTSINDLLFGIQAGPEQYSAWVNSLPGSPGLVDYTLEPLHVLLDSQDPRREALRRALSQYLTDRARWRDCS
EEKKKQHKMATSFHQTYRERHVEVLGGPLDSTHDLLFGNQATPEQFSTWTASLPSNPGLVDYSLEPLHTLLEEQNPKREALRQAISHYIMSRARWQNCS
VPSTRKYIYVGNFHQGFNERFREVAGGNAM------FDLHNHLSGENKWLKSTEKDPALLNYSLMPIHTLINSRSPKRESLRIALSEYIKKRSLKIEC-
KQSQDKKLSAHSFANSFSDRLTEITGGHTQ-DSALLFSASNDPGAYKQWLSSVPQQPDVISFSLKPLHMLMPAKNPKRDQLRRAIHDYVLQKALLKNCS
                  GC     RC      V   R                S          L          WW   W

  381       391       401       411       421       431       441         451       461       471
RPCPPGRQKSPRDPCQCVCHGSAVTTQDCCPRQRGLAQLEVTFIQAWGLWGDWFTATDAYVKLFFGGQEL--RTSTVWDNNNPIWSVRLDFGDVLLATG
RPCRSGQHKSSHDSCQCECQDSKVTNQDCCPRQRGLAHLVVSNFRAEHLWGDYTTATDAYLKVFFGGQEF--RTGVVWNNNNPRWTDKMDFENVLLSTG
-PCSGNTYPSQREDCSCSCSPSKYTNSDCCPTRKGLAKLIIEIRSATGLNGDYFSSTDSYVNFYFNGYEK--ETETIFDNDDPMWNETLDFGMVELGPL
SPCKTGIATNPKEPCACSCHNNPAVKANCCPAQRGIAQITVTVMRATGLFGDYFTWTDGYVKVFRNQKLFLGETPVIWNQNSPKWNWRFDLGSFVVSQF
              R                W          C    S  EY    M  C             M        L

     481       491       501       511       521       531       541       551
GPLRLQVWDQDSGRDDDLLGTCDQAPKSGSHEVRCNLNHGHLKFRYHARCLPHLGGGTCLDYVPQMLLGEPPGNRSGAVW-------------------
GPLRVQVWDADYGWDDDLLGSCDRSPHSGFHEVTCELNHGRVKFSYHAKCLPHLTGGTCLEYAPQGLLGDPPGNRSGAVW-------------------
KKYTIEVWDEDITTDNDLLLTCKGILKAGNNKMSCQYSDMQLVYTLYTTCAPYLGGEYCDEYIPSSP--------------------------------
GGLKLEVWDRDNKWDDDLLGTCNIQLKAGVKNDFCRLNHGLLYFKTSVTCGPSLAGSSCTEYVGSPMA-----SHLEKLYVSRHARPVPKDMLVKMGVL
     P    G    N                                 S

---------------------
---------------------
---------------------
LKERRALLGQTSEAESKRQNL
```

**Supplementary Figure 4. Sequence alignment of vertebrate perforins.**
Conserved residues mutated in FHL patients are highlighted in red, and non-conserved residues in brown.

Supplementary figure 5

A

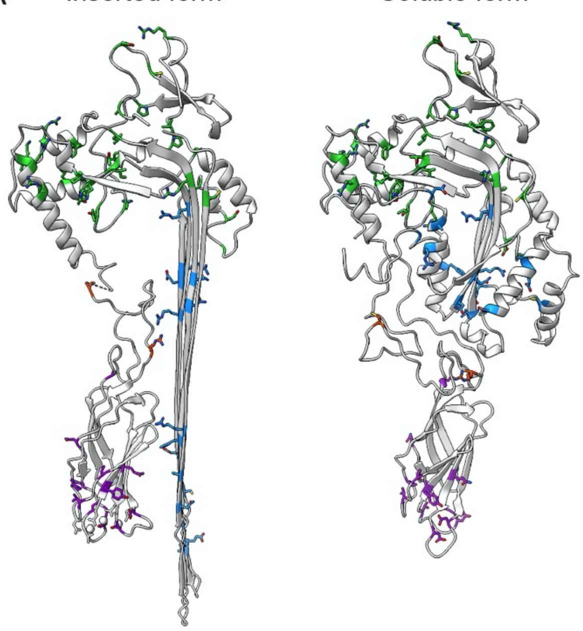

B

| | | | |
|---|---|---|---|
| M1V | R126C | R232H | C393R |
| R28C | G149S | R240G | R410P |
| V38M | F157V | N252S | R410W |
| V38L | S168N | E261K | F421C |
| P39H | S170R | A262D | G426S |
| G45E | R177C | C279Y | G429E |
| D49N | V183G | E298G | D430Y |
| V50M | F193L | R299C | T435M |
| R54C | G198R | G305R | Y438C |
| D70Y | P201T | G306C | T450M |
| C73R | L215I | D313V | T453M |
| P89T | G220S | G317R | P459L |
| A91V | T221I | G334S | Q481P |
| W95R | H222R | P345L | D486G |
| C102F | I224D | R356W | D491N |
| R119W | R225W | R357W | C525S |
| R123H | R232C | R361W | |

**Supplementary Figure 5. FHL-associated mutations of perforin identified in patients.**
A. Models of the inserted and soluble forms of perforin with identified mutations mapped on the structure as coloured sticks. Colour scheme is the same as in Figure 1. B. List of identified perforin mutations, with those considered to be benign polymorphisms underlined. All numbering is given for the human protein.
